# Transcriptomes of Electrophysiologically Recorded Dbx1-derived Inspiratory Neurons of the preBötzinger Complex in Neonatal Mice

**DOI:** 10.1101/2021.08.01.454659

**Authors:** Prajkta S. Kallurkar, Maria Cristina D. Picardo, Yae K. Sugimura, Margaret S. Saha, Gregory D. Conradi Smith, Christopher A. Del Negro

## Abstract

Breathing depends on interneurons in the preBötzinger complex (preBötC) derived from *Dbx1*-expressing precursors. Here we investigate whether rhythm- and pattern-generating functions reside in discrete classes of Dbx1 preBötC neurons. In a slice model of breathing with ∼5 s cycle period, putatively rhythmogenic Type-1 Dbx1 preBötC neurons activate 100-300 ms prior to Type-2 neurons, putatively specialized for output pattern, and 300-500 ms prior to the inspiratory motor output. We sequenced Type-1 and Type-2 transcriptomes and identified differential expression of 123 genes including ionotropic receptors (*Gria3* and *Gabra1*) that may explain their preinspiratory activation profiles and Ca^2+^ signaling (*Cracr2a, Sgk1*) involved in inspiratory and sigh bursts. Surprisingly, neuropeptide receptors that influence breathing (e.g., µ-opioid and bombesin-like peptide receptors) were only sparsely expressed, which suggests that cognate peptides and opioid drugs exert their profound effects on a small fraction of the preBötC core. These data in the public domain help explain breathing’s neural origins.

## Introduction

Inspiration, the preeminent active phase of breathing, originates in the preBötzinger complex (preBötC) of the lower brainstem^1,2^. Interneurons derived from *Dbx1*-expressing precursors (hereafter, Dbx1 neurons)^3,4^ comprise the preBötC core; they are responsible for generating inspiratory rhythm and transmitting it as a rudimentary output pattern to premotoneurons and motoneurons for pump and airway muscles^5–9^.

Cellular-level studies of inspiratory rhythm and pattern take advantage of transverse slices that retain the preBötC and remain spontaneously rhythmic *in vitro*. Constituent preBötC neurons can be recorded at the rostral slice surface while monitoring the inspiratory motor rhythm (∼5 s cycle period) via the hypoglossal (XII) cranial nerve. Inspiration begins with a low amplitude preinspiratory phase attributable solely to rhythmogenic neurons. As their activity crosses threshold, preinspiration leads to an inexorable high amplitude inspiratory burst, which recruits an additional class of pattern-related neurons that drive motor output^2,10–12^. There are two theories that differentiate the *rhythm* and *pattern* subsets of the Dbx1 preBötC neuron population.

The first theory posits that the neuropeptide somatostatin (SST) is a marker for output/pattern neurons. SST-expressing (SST^+^) preBötC neurons discharge during inspiration and postinspiration, i.e., the output phases of the inspiratory breathing cycle, rather than during the rhythmogenic preinspiratory phase. Furthermore, photostimulation in the preBötC of adult mice that express channelrhodopsin (ChR2) in SST^+^ neurons preferentially affects inspiratory motor pattern^6^. These findings, in the context of what we already know about Dbx1 neurons, suggest that the Dbx1 SST^+^ preBötC neurons play a dominant role in inspiratory pattern-generation and, by exclusion, that Dbx1 preBötC neurons lacking SST expression (SST^-^) are inspiratory rhythmogenic^6^.

The second theory subdivides Dbx1 preBötC neurons electrophysiologically. Neurons that activate with a ramp-like summation of synaptic potentials 300-500 ms before the onset of a large-magnitude inspiratory burst^13,14^ are considered “Type-1” and putatively rhythmogenic. Type-1 neurons also express A-type transient K^+^ current (*I*_A_) whose blockade perturbs preinspiratory activity and destabilizes the inspiratory rhythm *in vitro*^15^. Neurons that activate ∼300 ms later than Type-1^13,14^ are considered “Type-2”, putatively downstream from Type-1 and tasked with generating preBötC output^10,14^. Type-2 neurons express hyperpolarization-activated cationic current (*I*_h_)^13^ whose blockade profoundly affects motor output with mild effects on rhythm^16^.

We subdivided *rhythm* and *pattern* Dbx1 preBötC neurons based on the latter theory, which provides multiple criteria that can be measured during whole-cell recordings. We retrieved cytoplasmic contents and performed next-generation RNA sequencing on 17 samples: 7 Type-1, 9 Type-2, and 1 neuron, referred to here as Unknown, that did not precisely fit either category but was Dbx1-derived and inspiratory. These data elucidate the transcriptional profile at the cellular point of origin for breathing, a key physiological behavior for humans and all mammals. The data are publicly available (National Center for Biotechnology Information [NCBI] Gene Expression Omnibus [GEO] Accession code: GSE175642) to facilitate future studies of the Dbx1 preBötC core that interrogate the neural mechanisms of breathing.

## Results and Discussion

We analyzed Dbx1 preBötC neurons using Patch-Seq^17^, which entails whole-cell patch-clamp recording followed by next-generation sequencing (Supplementary Fig. 1A) and bioinformatics (Supplementary Fig. 1B).

The maximum yield of high-quality RNA was inversely proportional to whole-cell recording duration (5 min on average, 3-8 min in all experiments). Inspiratory burst characteristics and intrinsic membrane properties for Type-1 and Type-2 neurons (Fig. 1A top and bottom, respectively) are already well established^13,14^. Given the time constraints, we focused on measuring the intrinsic membrane properties i.e., delayed excitation and sag potentials, at the expense of recording fewer inspiratory burst cycles.

**Figure 1.**
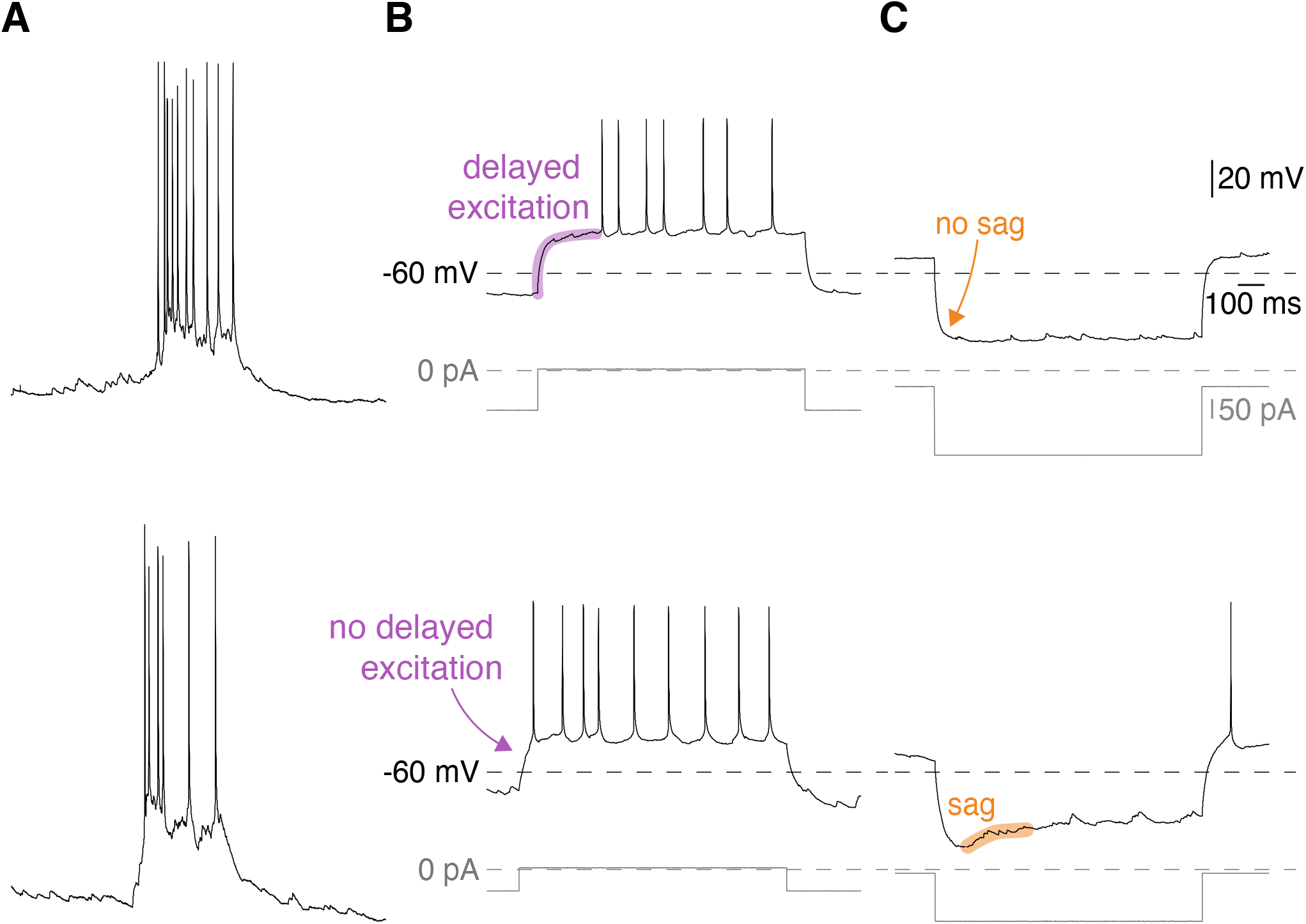
Physiological properties of Dbx1 preBötC inspiratory neurons. **A**, top and bottom traces show inspiratory burst characteristics in Type-1 and Type-2 Dbx1 preBötC neurons, respectively. **B**, Depolarizing current pulses (750-1000 ms) were applied from a membrane potential of -70 mV. Top and bottom traces show presence and absence of delayed excitation (purple) in Type-1 and Type-2 neurons, respectively. C, Hyperpolarizing current pulses (750-1000 ms) were injected from a membrane potential of -50 mV. Top and bottom traces show absence and presence of a sag potential (orange) in Type-1 and Type-2 neurons, respectively. Voltage, current, and time calibration bars apply to all traces.

Our patch-clamp recordings confirmed the previously published disparities between Type-1 and Type2 neurons. Type-1 neurons showed delayed excitation of 167 ± 40 ms (n = 7) when subjected to suprathreshold current steps from a baseline membrane potential below -70 mV (i.e., evidence of *I*_A_; Fig. 1B, top trace) but negligible sag potentials (2 ± 1 mV, n = 7) when subjected to hyperpolarizing current steps from a baseline membrane potential of -50 mV (i.e., no evidence of *I*_h_; Fig. 1C, top trace).

Type-2 neurons exhibited minimal delays in excitation (76 ± 40 ms, n = 9) when subjected to suprathreshold current steps from a baseline membrane potential below -70 mV (i.e., no evidence of *I*_A_; Fig. 1B, bottom trace) but their membrane potential trajectory ‘sagged’ 11 ± 4 mV (n = 9) in the direction of baseline when subjected to hyperpolarizing current steps from a baseline membrane potential of -50 mV (i.e., evidence of *I*_h_; Fig. 1C, bottom trace).

The disparities between delayed excitation and sag potentials measured in Type-1 and Type-2 neurons are unlikely to occur by random sampling from a single population with normally distributed expression of *I*_A_ and *I*_h_ with probabilities of p_delay_ = 0.0006 (t = 4.53, df = 13) and p_sag_ = 0.0002 (t = 4.96, df = 14), respectively. Therefore, we reject the null hypothesis and reconfirm that Type-1 and Type-2 are unique subpopulations of Dbx1 preBötC neurons^13^.

One Dbx1 preBötC inspiratory neuron we recorded and sequenced did not fit the criteria for Type-1 or Type-2, so we analyzed it as an Unknown.

We mapped all 17 samples to the murine genome (mm10 from *[Ensembl](http://nov2020.archive.ensembl.org/Mus_musculus/Info/Index)*); 83% ± 3% of the sequences aligned uniquely resulting in an average of 10,335,384 uniquely aligned reads (Supplementary Table 1).

### Transcriptomic differences between Type-1 and Type-2 neurons

The 31,543 genes that were expressed in at least one sample (7 Type-1 neurons and 9 Type-2 neurons) were examined for differential expression by DESeq2 (Fig. 2A). The Unknown neuron was not included in this analysis. DESeq2 identified 123 differentially expressed (DE) genes (Figs. 2A_a_, 3, Supplementary Table 2; p_adj_ < 0.01, log_2_ fold change [L2FC] > 1.5). The DESeq2 results were computed on Type-2 versus Type-1 neurons; a positive L2FC implies gene upregulation in Type-2 neurons, whereas a negative L2FC indicates gene upregulation in Type-1 neurons.

**Figure 2.**
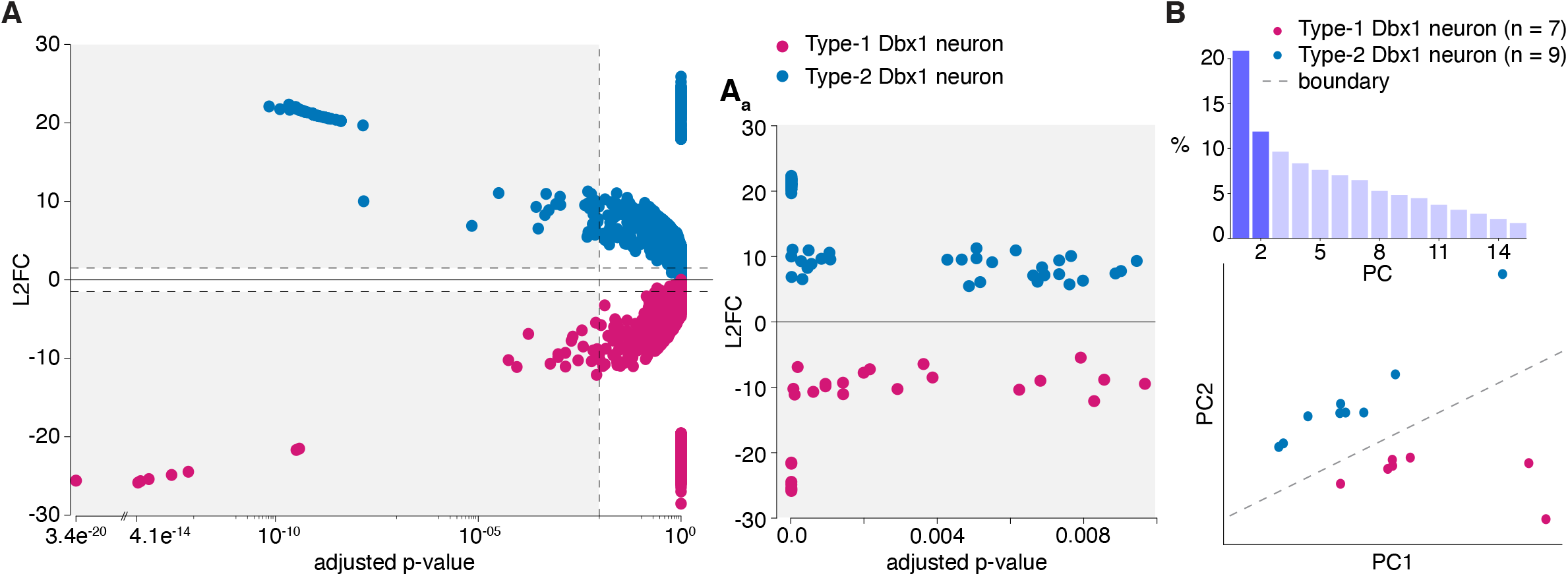
Transcriptomic differences in Type-1 and Type-2 neurons. **A**, L2FC versus adjusted p-values of 31,543 genes computed using DESeq2. Genes upregulated in Type-1 are in magenta and those upregulated in Type-2 are in blue-cyan. Gray shaded represents the region expanded in **A**_**a**_, which corresponds to L2FC > 1.5 and padj < 0.01. **B**, Bar chart (top) shows PCs (x-axis) and variability (y-axis) in the expression of 123 DE genes. Scatter plot (bottom) shows clustering of Type-1 (magenta) and Type-2 (blue-cyan) neurons using the first two PCs (highlighted in dark violet at top). The gray dashed line shows the boundary between the clusters of Type-1 and Type-2 neurons drawn by LDA.

We used principal component analysis (PCA) to assemble Dbx1 preBötC neurons in a plane based on transcriptome similarity, which revealed that Type-1 and Type-2 neurons form two distinct clusters (Fig. 2B). When we scrambled their Type-1 or Type-2 identities, PCA failed to differentiate the samples as two discrete classes (Supplementary Fig. 2), which suggests that Type-1 and Type-2 neurons are separate neuron classes based on their transcriptome (Fig. 2B and Supplementary Fig. 2) in addition to their unique neurophysiological properties (Fig. 1).

#### Genes upregulated in Type-1

We report an upregulation of the 5-HT_1D_ receptor gene, *Htr1d* (Fig. 3, Supplementary Fig. 3, Supplementary Table 2). *I*_A_, a characteristic Type-1 feature, is subject to neuromodulation by serotonin (5-HT) receptors, in mouse trigeminal ganglion neurons^18^ and CA1 pyramidal neurons^19,20^. Interestingly, in rhythmic slices from neonatal rats, bath application of 5-HT increases the frequency, but not the amplitude, of XII motor output, implying a selective effect on the rhythm-generating population^21–23^, which probably maps to Type-1 neurons.

**Figure 3.**
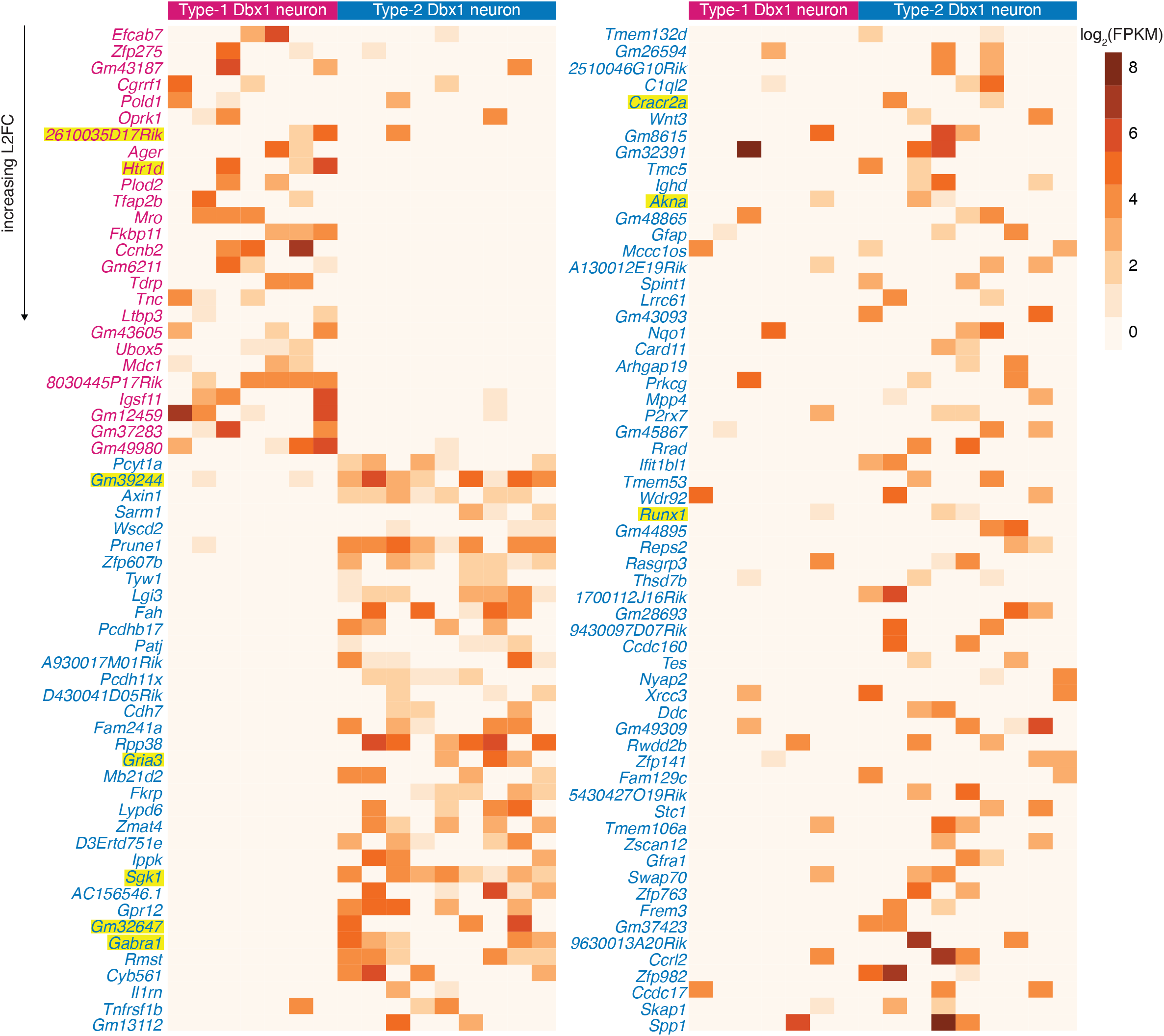
Log_2_(FPKM) of the DE genes in Type-1 and Type-2 neurons. Log_2_(FPKM) value is indicated by a pseudocolor scale. Individual neurons are listed in columns. Genes are listed in rows, arranged in the increasing order of L2FC values. The names of genes upregulated in Type-1 are in magenta and those upregulated in Type-2 neurons are in blue-cyan. Genes names highlighted in yellow are mentioned in the text.

#### Genes upregulated in Type-2

Depletion of Ca^2+^ in the endoplasmic reticulum (ER) activates the stromal interaction molecule (STIM) proteins, which subsequently activate the Ca^2+^ release-activated Ca^2+^ (CRAC) channels on the plasma membrane via the key subunit of CRAC channel, Orai1^24–27^. We report Type-2 upregulation of the CRAC channel regulator 2A gene, *Cracr2a*, as well as the serine/threonine protein kinase gene, *Sgk1*, which activates STIM1 and Orai1 and thus enhances store-operated Ca^2+^ entry (SOCE)^28^. SOCE-related mechanisms that could support or augment inspiratory (eupnea-related) and sigh-related pattern generation remain important topics for investigation. For example, regarding inspiration, intracellular Ca^2+^ signaling in the context of SOCE could recruit Ca^2+^-activated non-specific cationic current (*I*_CAN_), which profoundly contributes to inspiratory burst pattern^29–31^, consistent with the role of Type-2 neurons. Sigh breaths, which occur at lower frequencies but are two-fold larger in magnitude^32^ are likely also to involve Ca^2+^ signaling and possibly SOCE mechanisms that recruit *I*_CAN_^33^.

Dbx1 preBötC neurons are glutamatergic^3,4^ and excitatory synaptic interactions, predominantly mediated by postsynaptic AMPA receptors, are essential for inspiratory rhythm and pattern generation^34,35^. We detect Type-2 upregulation of the AMPA receptor, *Gria3* (Figs. 3, 4, Supplementary Table 2), which may at first seem counterintuitive for the neuron class with shorter inspiratory drive latency and typically lower-amplitude inspiratory bursts^10,13,14^. Because the longer inspiratory drive latency in Type-1 neurons may be attributable to the rich topology of their excitatory synaptic interconnections^36,37^, we surmise that upregulation of *Gria3* in Type-2 neurons augments inspiratory drive in these less richly interconnected preBötC neurons. Upregulation of *Gria3* may amplify postsynaptic AMPA receptor-mediated inspiratory drive to accomplish the Type-2 role as output neurons.

**Figure 4.**
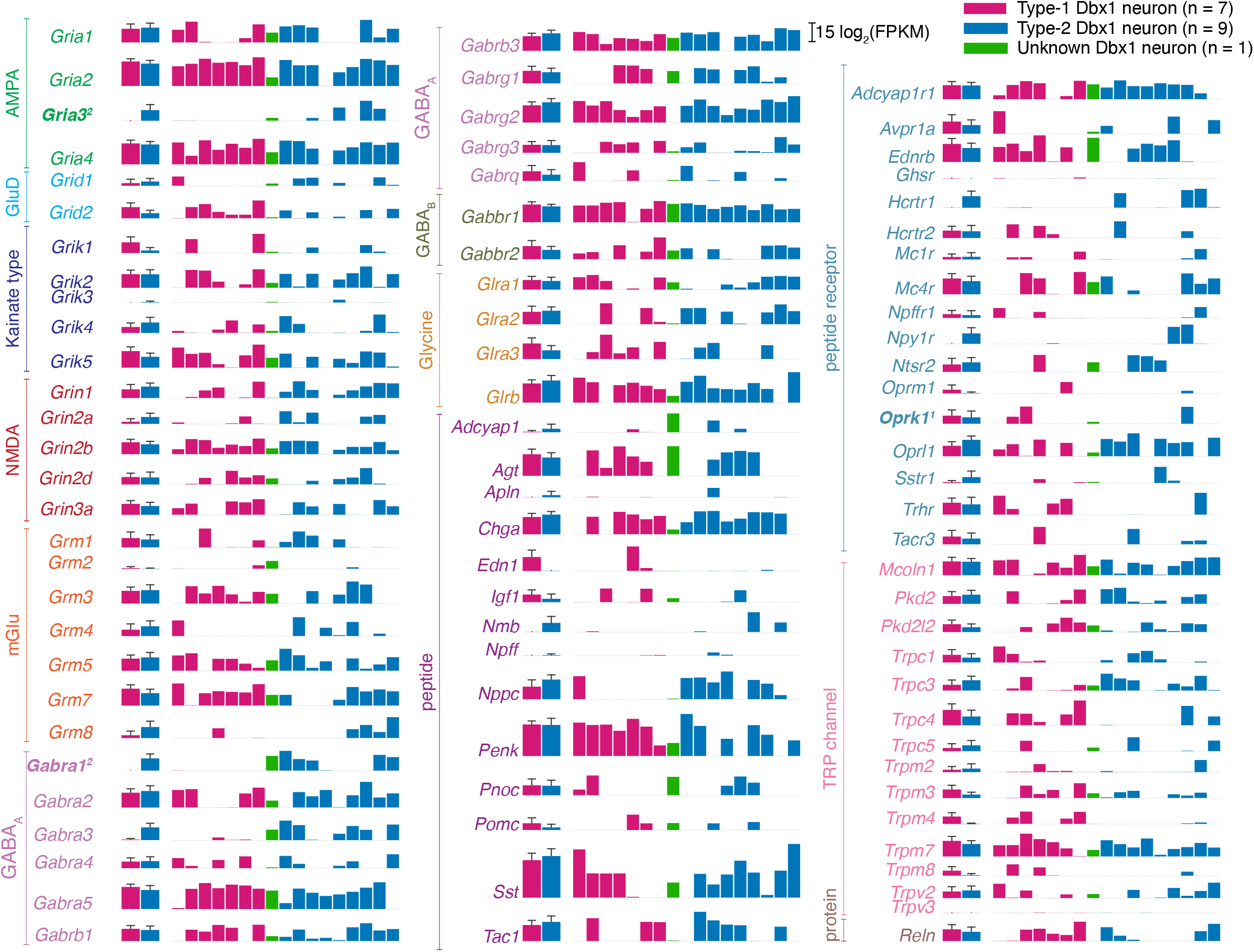
Quantitative gene expression for ionotropic and metabotropic synaptic receptors, neuropeptides, neuropeptide receptors, and Trp channels. The first two bars show group data for Type-1 (n = 7; magenta bar) and Type-2 (n = 9; blue-cyan bar). The height of the bar is log_2_(mean FPKM) value and the error bar with horizontal cap shows log_2_(mean + SD). The next set of 17 bars shows log_2_(FPKM) values for each neuron in the following order: 7 Type-1 neurons (magenta), 1 Unknown neuron (green), and 9 Type-2 neurons (blue-cyan). Gene names are color-coded according to the subfamily to which they belong. Gene names in bold indicate DE and contain a superscript 1 if upregulated Type-1 neurons and 2 if upregulated in Type-2 neurons.

Synaptic inhibition influences inspiratory rhythm and output pattern^5,38–41^. We report upregulation of the GABA_A_ receptor, *Gabra1*, in Type-2 neurons (Figs. 3, 4, Supplementary Table 2). GABAergic drive regulates Type-1 neurons, but *Gabra1* upregulation in Type-2 neurons suggests that inhibitory inputs may be equipped to bypass the oscillator and selectively act on the pattern-generating subpopulation to arrest inspiration with immediacy. Behavioral exigencies that might apply include the breath-hold dive reflex upon submersion or attentive immobility, that is, the arrest of all movement (including breathing) for predators stalking prey or prey attempting to camouflage themselves in the context of being hunted.

Transcription factors program cell fate during embryonic development and regulate gene expression postnatally. Because this study was performed postnatally (P0-2) it cannot detect the transcription factors acting in precursor cells. Nevertheless, we report upregulation of transcription factors *Akna* and *Runx1* in Type-2 neurons (Fig. 3, Supplementary Table 2). *Runx1* helps consolidate spinal motor neuron identity by suppressing interneuron programs^42^. It may, therefore, seem counterintuitive that *Runx1* is upregulated in Type-2, but we speculate that it may be acting to suppress Type-1 programs or else halting any further programming or developmental changes to Type-2 neurons. The potential role of *Akna* is not known.

#### Non-coding RNA

We report 15 differentially expressed long non-coding RNA (such as *2610035D17Rik, Gm39244, Gm32647*) (Fig. 3, Supplementary Table 2). The function of these transcripts and their role(s) in inspiratory rhythm- and pattern-generation is unexplored for now.

### Transcripts associated with cellular neurophysiology

We next examined a broad spectrum of ionotropic and metabotropic synaptic receptors, peptides, peptide receptors, and transient receptor potential (Trp) ion channels (Fig. 4); voltage-dependent ion channels, regulatory subunits, and intracellular receptors (Fig. 5); purine receptors, monoamine receptors, and cell adhesion molecules (Supplementary Fig. 3); as well as transcription factors (Supplementary Fig. 4) irrespective of whether they are DE or non-DE genes. Here, our goal was to understand preBötC neuron excitability and signaling in general, not differential expression, so the criteria for inclusion were relaxed: any genes that were expressed in >25% of the neurons (4 out of 17), regardless of the type of neuron, were quantified (Supplementary Table 3) and cataloged.

**Figure 5.**
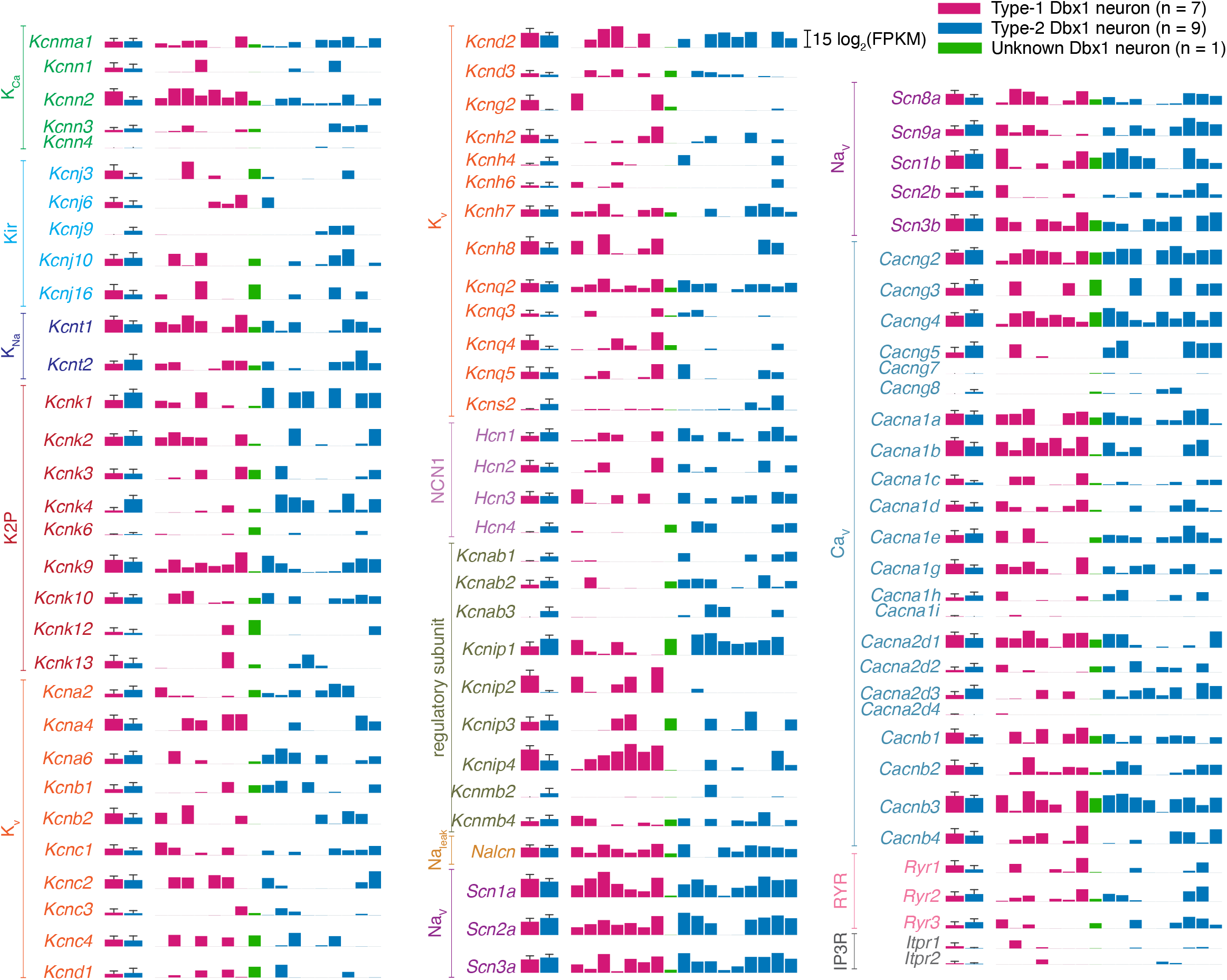
Quantitative gene expression for voltage-dependent ion channels, regulatory subunits, and intracellular receptors. The first two bars show group data for Type-1 (n = 7; magenta bar) and Type-2 (n = 9; blue-cyan bar). The height of the bar is log_2_(mean FPKM) value and the error bar with horizontal cap shows log_2_(mean + SD). The next of 17 bars shows log_2_(FPKM) values for each neuron in the following order: 7 Type-1 neurons (magenta), 1 Unknown neuron (green), and 9 Type-2 neurons (blue-cyan). Gene names are color-coded according to the subfamily to which they belong. None are DE.

#### Ion channels and their regulation

The delayed excitation in Type-1 neurons is a functional characteristic of A-type transient K^+^ current that can be mediated by the K_v_ channels: K_v_1.4 (*Kcna4*), K_v_3.3-3.4 (*Kcnc3-4*), and K_v_4.1-4.3 (*Kcnd1-3*). However, the magnitude of *I*_A_ depends on two proteases (*Dpp6, Dpp10;* Supplementary Table 3) that increase the plasma membrane expression of K_v_4.2^43,44^ and the interacting proteins for all K_V_4.x channels (K channel-interacting proteins i.e. KChIPs [*Kcnip1-4*]; Fig. 5, Supplementary Table 3) that substantially enhance *I*_A_ without affecting its voltage-dependence or kinetics^45,46^. Neither the genes for K_v_ ion channels nor these regulatory genes cross our significance threshold in DESeq2 so we make no statistical claims; nevertheless, we posit that *Kcnip2* and *Kcnip4* are important for *I*_A_ in Type-1 but our methods and analyses were insufficient to detect them.

The sag potential in Type-2 neurons is a classic property of hyperpolarization-activated, cyclic nucleotide-gate channels (HCN) channels encoded by *Hcn1-4* (Fig. 5, Supplementary Table 3). We do not detect any significant difference in the expression of these genes.

Our data show that Type-1 and Type-2 neurons comprise electrophysiologically discrete classes (Fig. 1) and yet they do not express different levels of ion channels typically associated with *I*_A_ and *I*_h_ (Fig. 5). We propose that the magnitude of *I*_A_ and *I*_h_, and thus their influence on membrane potential trajectories, are instead governed by neuromodulation, upstream influences that would be difficult to assess by Patch-Seq, the potential role of *Htr1d* notwithstanding (see above).

#### Peptides and peptide receptors

Given the importance of the µ-opioid receptor, *Oprm1*, in opioid-induced respiratory depression we analyzed the expression of *Oprm1*, even though it was neither DE nor expressed in >25% of the samples. *Oprm1* is expressed in only 3 out 17 samples here (2 Type-1 neurons and 1 Type-2 neuron express *Oprm1*, net mean expression of 0.50 ± 1.58 log_2_[FPKM], n = 17; Fig. 4, Supplementary Table 3). Our results are congruent with the recent demonstration that ∼8% of Dbx1 preBötC neurons express *Oprm1*^47^. The apparent sparsity of *Oprm1* expression does not negate the obvious potency of µ-opioid receptor-mediated effects in the preBötC but it constrains the mechanism underlying opioid-induced respiratory depression to operate on a small fraction of constituent preBötC neurons, both Type-1 and Type-2, which affect rhythm and pattern.

Peptide receptors for tachykinins (neurokinin-1 receptor specifically), neuromedin B, and gastrin-releasing peptide (NK1R, Nmbr, and Grpr, respectively) modulate breathing by acting directly on preBötC neurons. NK1R-expressing (*Tacr1-3*) preBötC neurons form a heterogeneous population of rhythm- and pattern-generators^48^. Consistent with this idea, we do not observe differential expression of genes encoding NK1R (Fig. 4, only *Tacr3* passed our screening criteria for display).

Sigh breaths draw on the inspiratory reserve volume of the lungs to reinflate collapsed (or collapsing) alveoli and optimize pulmonary function^32^. Bombesin-like peptide receptors, Nmbr and Grpr, that modulate sighing were detected in ∼7% of all preBötC neurons, including Dbx1 and non-Dbx1 subpopulations^49^. We report more than double that expression level, i.e., ∼18% of our samples (n = 3) expressed bombesin-like peptide receptor transcripts: 1 Type-1 and 1 Type-2 neuron expressed *Nmbr* for a combined mean expression level of 0.05 ± 0.19 log_2_(FPKM) and 1 Type-2 neuron expressed *Grpr* at an expression level of 0.01 log_2_(FPKM). These data are in line with expectations because our study focuses exclusively on Dbx1 preBötC neurons, the core for inspiratory breathing rhythm and pattern, whereas Li *et al*. measured transcripts in any preBötC neuron, ∼50% of which are not derived from Dbx1-expressing progenitors.

Pituitary adenylate cyclase-activating peptide (PACAP) is important for breathing responses to CO_2_. PACAP mutant mice exhibit blunted chemosensitivity and die within 3-weeks after birth due to respiratory defects^50^. Bilateral microinjections of PACAP in preBötC, *in vivo*, increases breathing frequency and inspiratory motor output^50^ and *in vitro* leads to an increase in XII motor output frequency^50,51^. We report expression of PACAP receptor (*Adcyap1r1*) in 15 out of 17 samples (6 Type-1 neurons, 8 Type-2 neurons and 1 Unknown) for a combined mean expression level of 3.52 ± 3.24 log_2_(FPKM), which would explain the effect of PACAP on both, rhythm and pattern (Fig. 4, Supplementary Table 3).

SST (and SST receptors) are expressed in core preBötC neurons, including – but not limited to – those derived from Dbx1-expressing precursors^3,4,52^. Several seminal studies *in vitro* and *in vivo* show that SST modulates inspiratory rhythm and pattern^53^. However, more recent studies specifically posit that SST-expressing (SST^+^) preBötC neurons are inspiratory pattern generators, i.e., output-related neurons^6,10^. We report *Sst* expression in 14 out of 17 Dbx1 neurons: 5 Type-1, 8 Type-2, and 1 Unknown for a combined mean expression level of 10.11 ± 11.65 log_2_(FPKM) (Fig. 4). Therefore, our data are incongruent with the theory that divides *rhythm* and *pattern* Dbx1 preBötC neurons on the basis of SST. Since we find that SST^+^ Dbx1 neurons are well apportioned among putative rhythm- and pattern-generating subpopulations, Type-1 and Type-2, respectively, we posit that a substantial population of non-Dbx1 SST^+^ neurons with exclusively pattern-generating functionality exists in the preBötC, which we did not sample.

### Perspectives

Dbx1-derived preBötC neurons operate in unison to generate breathing movements. We distinguish *rhythm* and *pattern* functions in seeking to understand the neural origins of breathing but we should remember that there is just one behavior. Type-1 and Type-2 neurons are significantly different on the basis of only 123 genes out of the 31,420 genes, although it should be noted that our criteria of p_adj_ < 0.01 and L2FC > 1.5 are relatively stringent^54–56^. Thus, these putative rhythm- and pattern-generating populations have much more in common in terms of their transcriptomes, compared to that which differentiates them.

We acknowledge technical limitations could have limited our ability to resolve Type-1 vs. Type-2 transcriptome disparity. The minute amount of starting material (usually ≤10 pg RNA in the retrieved cytoplasm) from a single neuron necessitates substantial amplification to get sufficient cDNA for sequencing (>150 pg/µL). Amplification leads to bias favoring some sequences and invariably drowning-out others^57^. Also, reverse transcription (RT) errors can leads to misreplication follow by a failure to map to the reference genome. These caveats produce false zeros for genes that are biologically expressed at non-zero levels^54,58,59^. We performed multiple quality checks (Supplementary Fig. 1B) for our sequences, used stringent criteria for selecting DE genes and scrambling to ensure the differential expression analysis was efficient in detecting DE genes. However, these checks cannot differentiate a technical zero from a biological zero. The upshot of these caveats is that our Patch-Seq analysis assuredly missed some expressed genes and incorporates some false zeros. We further acknowledge that the disparities between Type-1 and Type-2 neurons may come about during translation and post-translational modifications, which impact phenotypic properties like inspiratory burst magnitude (*I*_CAN_, i.e., Trpm4-dependent), delayed excitation (*I*_A_), and sag potentials (*I*_h_).

Nevertheless, these data provide the electrophysiology and transcriptomic data, including non-coding transcripts, for Dbx1 preBötC inspiratory neurons at the core of inspiratory rhythm and pattern generation. The transcriptomic data reported here can be utilized or meta-analyzed to design new experiments studying the neural control of breathing.

## Methods

The Institutional Animal Care and Use Committee at William & Mary approved these protocols, which conform to policies set forth by the Office of Laboratory Animal Welfare (National Institutes of Health) and the National Research Council^60^.

### Mice

We crossed knock-in mice generated by inserting an *IRES-CRE-pGK-Hygro* cassette in the 3’ UTR of the *Developing brain homeobox 1* (i.e., *Dbx1*) gene, i.e., Dbx1^Cre^ mice^61^ (IMSR Cat# EM:01924, RRID:IMSR_EM:01924) with mice featuring Cre-dependent expression of fluorescent Ca^2+^ indicator GCaMP6f, dubbed Ai148 by the Allen Institute (RRID: IMSR_JAX:030328, Daigle et al. 2018). Ai148 mice had C57Bl/6J background; Dbx1^Cre^ mice had a mixed C57Bl/6J;CD1 genetic background. We used their offspring, referred to as Dbx1;Ai148 mice (P0-2) for experiments.

The animals were housed in colony cages on a 12/12 h light/dark cycle with controlled humidity and temperature at 23 ºC and fed *ad libitum* on a standard commercial mouse diet (Teklad Global Diets, Envigo) with free access to water.

### *In vitro* slice preparations

The workbench was cleaned with RNase ZAP (Thermo Fisher, Waltham, MA) before beginning each experiment. All the beakers and tools were either autoclaved or cleaned first with RNase ZAP and then with nuclease-free water (NFW).

Dbx1;Ai148 pups were anesthetized by hypothermia, consistent with the American Veterinary Medical Association (AVMA) guidelines for euthanasia^63^. The neuraxis, from the pons to lower cervical spinal cord, was removed within ∼2 min and submerged in ice-cold artificial cerebrospinal fluid (aCSF) containing (in mM): 124 NaCl, 3 KCl, 1.5 CaCl_2_, 1 MgSO_4_, 25 NaHCO_3_, 0.5 NaH_2_PO_4_, and 30 dextrose. The aCSF was prepared in an RNase-free environment and then aerated continually with 95% O_2_ and 5% CO_2_ during the experiment. We trimmed the neuraxis and glued the dorsal surface of the brainstem onto an agar block (exposing the ventral side). The block and brainstem were affixed rostral side up within a vibratome (Campden Instruments 7000 smz-2, Leicester, UK) while perfusing with aerated ice-cold aCSF. We cut a single transverse slice 450-500 µm thick with preBötC on its rostral surface. We started a 3-hr countdown clock as soon as the mouse was anesthetized and discarded the slice at the end of the interval to avoid sample degradation and contamination.

### Whole-cell patch-clamp recording and cytoplasmic sample collection

We perfused slices with aCSF (28 °C) at 2-4 ml/min in a recording chamber on a fixed-stage upright microscope. The external K^+^ concentration ([K^+^]_ext_) of the aCSF was raised from 3 to 9 mM to facilitate robust respiratory rhythm and XII motor output. We recorded XII motor output using suction electrodes fabricated from autoclaved borosilicate glass pipettes (OD: 1.2 mm, ID: 0.68 mm) fire polished to a diameter of ∼100 µm. XII motor output was amplified by 2,000, band-pass filtered at 0.3-1 kHz, and RMS smoothed for display.

Inspiratory Dbx1 preBötC neurons were selected visually based on rhythmic fluorescence emitted by GCaMP6f. Patch pipettes were fabricated from autoclaved borosilicate glass (OD: 1.5 mm, ID: 0.86 mm) using a 4-stage program on a Flaming-Brown P-97 micropipette puller (Sutter Instruments, Novato, CA). Patch pipettes with tip resistance of 3-5 MΩ were filled with an internal solution, mixed in an RNase-free environment, containing (in mM): 123 K-gluconate, 12 KCl, 10 HEPES, 0.2 EGTA, 4 Mg-ATP, 0.3 Na-GTP, 10 Na_2_-phosphocreatine, and 13 Glycogen (osmolarity adjusted to 270-290 mOsm and pH 7.25). We added 0.8% Recombinant Ribonuclease Inhibitor (RRI) to the internal solution immediately before each experiment to preserve RNA. We used robotic micromanipulators (Sensapex, Helsinki, Finland) to guide our patch pipettes toward inspiratory neurons under visual control and then performed whole-cell patch-clamp recordings using an EPC-10 patch-clamp amplifier (HEKA Instruments, Holliston, MA) with PATCHMASTER software (RRID:SCR_000034).

Starting from a quiescent membrane potential between inspiratory bursts, we defined inspiratory drive latency as the elapsed time from first detection of summating synaptic potentials (EPSPs) until the onset of the inspiratory burst.

We tested for A-type K^+^ current (*I*_A_) by applying suprathreshold depolarizing current step commands of 750-1000 ms duration from holding potentials of -70 mV (Fig. 1B) and -50 mV. The net applied current during the step command was equivalent regardless of holding potential. Neurons expressing *I*_A_ exhibited delayed excitation of 120-220 ms from a holding potential of -70 mV, but not from a holding potential of -50 mV^13,14^.

We tested for hyperpolarization-activated cationic current (*I*_h_) by applying hyperpolarizing current step commands of 750-1000 ms duration, which caused initial voltage excursions exceeding -30 mV from a holding potential of -50 mV (Fig. 1C). Neurons expressing *I*_h_ exhibited a time- and voltage-dependent depolarizing ‘sag’^13,14^.

After, categorizing the Dbx1 preBötC neuron as Type-1 and Type-2 (Supplementary Fig. 1A_a_), cytoplasmic contents were extracted under voltage clamp (−60 mV holding potential) by applying negative pressure (0.7-1.5 psi). Successful extraction left the neurons visibly shrunken. Negative pressure was applied for a maximum of 10 mins or until the neuron was electrophysiologically unstable, indicated by holding currents exceeding -600 pA, whichever happened first. The patch pipettes were retracted promptly, and the cytoplasmic contents were ejected by breaking the pipette tip at the bottom of the RNase-free PCR tube containing 4 µL of stock solution (stock solution = NFW with 2% RRI) while applying positive pressure (Supplementary Fig. 1Ab). Great care was taken to avoid any bubbles while applying positive pressure. Samples were briefly spun in a mini centrifuge, then snap-frozen in liquid nitrogen and stored at -80°C until further processing.

We monitored for potential contamination by collecting negative control samples during each experiment. Patch pipettes were filled with internal solution and then inserted into the tissue without targeting any neuron for recording; their contents were processed identically.

### cDNA synthesis, library preparation and sequencing

RNA from the recovered cytoplasm of patch-clamped neurons was converted to complementary DNA (cDNA) according to the SMART-Seq HT protocol (Takara Bio USA, Mountain View, CA), which incorporates the template-switching activity of the reverse transcriptase to select for full-length cDNAs and to add PCR adapters to both ends of the first-strand DNA (SMART = Switching Mechanism at 5’ end of RNA Template). Samples were denatured at 72 ºC for 3 min. poly(A)+ RNA was reverse transcribed using a tailed oligo(dT) primer. First strand cDNA synthesis and double-stranded cDNA amplification were performed in a thermocycler using the following program: 42 °C for 90 min; 95 °C for 1 min; 18 cycles of 98 °C for 10 s, 65 °C for 30 s, 68 °C for 3 min; and finally, 72 °C for 10 min. PCR-amplified cDNA was purified by immobilization on Agencourt AMPure XP beads (Beckman Coulter, Brea, CA), and then washed with 80% ethanol and eluted with elution buffer. Sequencing libraries were prepared from the amplified cDNA using SMART-Seq PLUS kits (Takara Bio USA, Mountain View, CA). Unique dual indexes were used on the amplified libraries to identify samples. We verified average cDNA size, abundance, and quality control of the final library using a Bioanalyzer High Sensitivity kit (Agilent, Santa Clara, CA) and Qubit dsDNA High-sensitivity Assay Kit (Molecular Probes, Eugene, OR) (Supplementary Fig. 1A_c_). cDNA samples containing less than 150 pg/µl cDNA were not sequenced. The cDNA sequencing libraries passing quality control were sequenced using an Illumina HiSeq X Sequencing system (Supplementary Fig. 1A_d_) with paired-end (150 bp) reads (Admera Health Biopharma Services, South Plainfield, NJ). A total of 18 samples were sequenced. Investigators were blinded to cell type during library construction and sequencing.

### Quality control, pre-processing, and alignment to reference genome

Nucleotide sequences along with their corresponding quality scores were returned as FASTQ files. We received an average of 18,724,864 (n = 18 samples) paired-end reads per sample. The quality of reads was verified using FastQC v0.11.8 (Supplementary Fig. 1B_a_). One sample returning 688 reads was discarded.

The mouse reference genome, mm10 from *[Ensembl](http://nov2020.archive.ensembl.org/Mus_musculus/Info/Index)*, was used to create the genome directory for aligning the reads in STAR using the following commands:

1. wget ftp://ftp.ensembl.org/pub/release-102/fasta/mus_musculus/dna/Mus_musculus.GRCm38.dna.primary_assembly.fa.gz
2. wget ftp://ftp.ensembl.org/pub/release-102/gtf/mus_musculus/Mus_musculus.GRCm38.102.gtf.gz
3. gunzip Mus_musculus.GRCm38.dna.primary_assembly.fa.gz
4. gunzip Mus_musculus.GRCm38.102.gtf.gz
5. STAR --runMode genomeGenerate --genomeDir {path to genome folder} --genomeFastaFiles Mus_musculus.GRCm38.dna.primary_assembly.fa --sjdbGTFfile Mus_musculus.GRCm38.102.gtf --sjdbOverhang 149 --genomeSAsparseD 2

The raw reads were aligned to the mm10 reference genome by the splice-aware STAR software v2.7.7a, which generates BAM alignment files (Supplementary Fig. 1B_c_), using the following command:

1. STAR --genomeDir mm10ReferenceGenome --readFilesIn inputFASTQFile1.fastq inputFASTQFile2.fastq --outFileNamePrefix outputBAMFile --outSAMtype BAM SortedByCoordinate --outReadsUnmapped Fastx

The alignment procedure (above) was done a total of 3 times for each sample to monitor the quality of the samples. The first alignment corresponds to raw reads. The second and third alignments are done after removing adapter and overrepresented sequences, respectively.

Adapters added during library construction: AGATCGGAAGAGCACACGTCTGAACTCCAGTCA (paired end 1) and AGATCGGAAGAGCGTCGTGTAGGGAAAGAGTGT (paired end 2) were trimmed (Supplementary Fig. 1B_b_) by bbduk v38.00 using the following command:

1. sh bbduk.sh in1=inputFASTQFile1.fastq. in2=inputFASTQFile2.fastq out1=outputFASTQFile1.fastq out2=outputFASTQFile2.fastq ktrim=r -Xmx27g mm=f k=33 hdist=1 literal=AGATCGGAAGAGCACACGTCTGAACTCCAGTCA,AGATCGGAAGAGCGTCGTGTAGGGAAAGAGT GT tbo tpe

The SMART-Seq HT kit uses dT priming for first-strand cDNA synthesis, annealing to the poly A tails of mRNA. The sequencing reads contained poly T/A sequences that were identified by FASTQC and tagged as overrepresented sequences, and finally trimmed (Supplementary Fig. 1B_b_) by cutAdapt v3.2 using the following command:

1. cutadapt -a overrepresentedSequence -A overrepresentedSequence’ -o outputFASTQFile1.fastq -p outputFASTQFile2.fastq inputFASTQFile1.fastq inputFASTQFile2.fastq -m 10 -j 4

The STAR v2.7.7a alignment software tallies the number of sequences that (*i*) align uniquely, (*ii*) align at multiple portions, or (*iii*) fail to align with mm10. We present these alignment statistics for each step (raw reads, adapter-trimmed reads, and adapter-trimmed reads following removal of overrepresented sequences, i.e., *processed reads*) in Supplementary Table 1. Only the final processed reads were used for downstream analysis.

We employed Qualimap v.2.2.2 to perform a final quality check of the BAM alignment files using this command:

1. qualimap rnaseq -bam inputFile.bam -gtf Mus_musculus.GRCm38.102.gtf outdir outputFileDir --paired --java-mem-size=4G

Intergenic reads exceeding 30% indicate DNA contamination. Our samples showed an average of 5.37% ± 2.33% intergenic reads (n = 17) so we conclude that our samples were not contaminated and thus could be used for downstream analyses. The quality control metrics of the processed reads, computed by Qualimap, are shown in Supplementary Table 1.

Uniquely aligned reads were converted to fragment counts (Supplementary Fig. 1B_d_) using featureCounts from the Rsubread package v2.4.2. The data pre-processing was performed using computing facilities at William & Mary (https://www.wm.edu/it/rc).

## Data analysis

We wrote custom R scripts (R v4.0.3, RRID:SCR_001905) that quantify gene expression as fragment counts per kilobase of exon per million mapped reads (FPKM; Supplementary Fig. 1B_e_). This quantification method is ideally suited for paired-end reads and normalizes for gene length and quantity of mapped reads. We also used R scripts to compute the mean and standard deviation (SD) of FPKM values. The log_2_ transformed value of FPKM, mean FPKM, or mean + SD FPKM were used for visualization.

Genes that are part of mm10 but had zero fragment counts in all 17 samples were omitted from all further analyses and consideration (23,824 genes). We performed differential expression analyses on the remaining non-zero genes (31,543 genes) using DESeq2 v1.30.1 software. DESeq2 uses fragment count (not FPKM) for each gene to calculate its geometric mean (non-zero counts only) across all the samples (Supplementary Fig. 1B_f_). Next, it normalizes each count by dividing the fragment count of the gene by its geometric mean. The fold change (L2FC) between Type-1 and Type-2 Dbx1 neurons is calculated using logarithm (base 2) of the normalized counts. Any gene where the L2FC exceeds 1.5 and adjusted p-value (p_adj_) is less than 0.01 was deemed differentially expressed (DE).

Custom MATLAB scripts (RRID:SCR_001622) implemented unsupervised principal component analysis (PCA) for dimensionality reduction and clustering of the log_2_(FPKM) expression profiles of 123 DE genes and 16 samples. Although the PCA was performed without regard for sample category, clustering of the Type-1 and Type-2 samples is evident using the first two principal component scores that represent 32% of the variation in the data (Fig. 2B, axes labelled PC1 and PC2). A boundary line calculated using linear discriminant analysis (LDA) shows that an accurate Type-1 vs. Type-2 classification may be performed using the first two principal component scores.

As a control, we permuted the labels identifying the 16 samples as Type-1 vs. Type-2 and repeated our analyses of the log_2_(FPKM) expression profiles. In each of 10 scrambled data sets, 50% of the samples were correctly labelled and 50% were “imposters” with false identities. In each case, we performed DESeq2 analysis to obtain a list of “DE genes” and performed PCA on this subset of genes. Supplementary Fig. 2A shows a representative LDA using PC1 and PC2 for genes differentially expressed between two groups of neurons with scrambled Type-1 and -2 identities. The classification error in this case is 0.18, because 3 of the 16 neurons are misclassified (3 apparent Type-2 neurons, one of which is a Type-2 imposter, are on or below the boundary line). Overall, the performance of classifiers obtained by LDA of such “DE genes” was severely degraded compared to the unscrambled case, especially when the LDA was restricted to the first several principal component scores (Supplementary Fig. 2). This result adds confidence to our list of DE genes for Type-1 versus Type-2 neurons.

## Data & code availability

The raw data of nucleotide sequences along with their corresponding quality scores (FASTQ format), raw fragment counts of the processed data (text file) and the FPKM values of the processed data (text file) are publicly available in the NCBI GEO database (Accession code: GSE175642). The custom R scripts written to process the raw fragments counts are freely available (https://github.com/prajkta9/bioinformatics-scRNA-seq).

## Acknowledgements

funded by NIH R01 NS107296 (PI: Del Negro), NIH R01 AT01816 (PI: Conradi Smith and Del Negro, NSF DMS 1951646 (PI: Conradi Smith), NSF DEB 2031275 (PI: Conradi Smith), NIH R15 HD096415 (PI: Saha)

## Notes

<u>Conflict of interest</u>: The authors declare no conflicting financial interests.

https://www.ncbi.nlm.nih.gov/geo/query/acc.cgi?acc=GSE175642

### Competing Interest Statement

The authors have declared no competing interest.

