## Supplementary Fig. 1 for "Transcriptomes of Electrophysiologically Recorded Dbx1-derived Inspiratory Neurons of the preBötzinger Complex in Neonatal Mice"

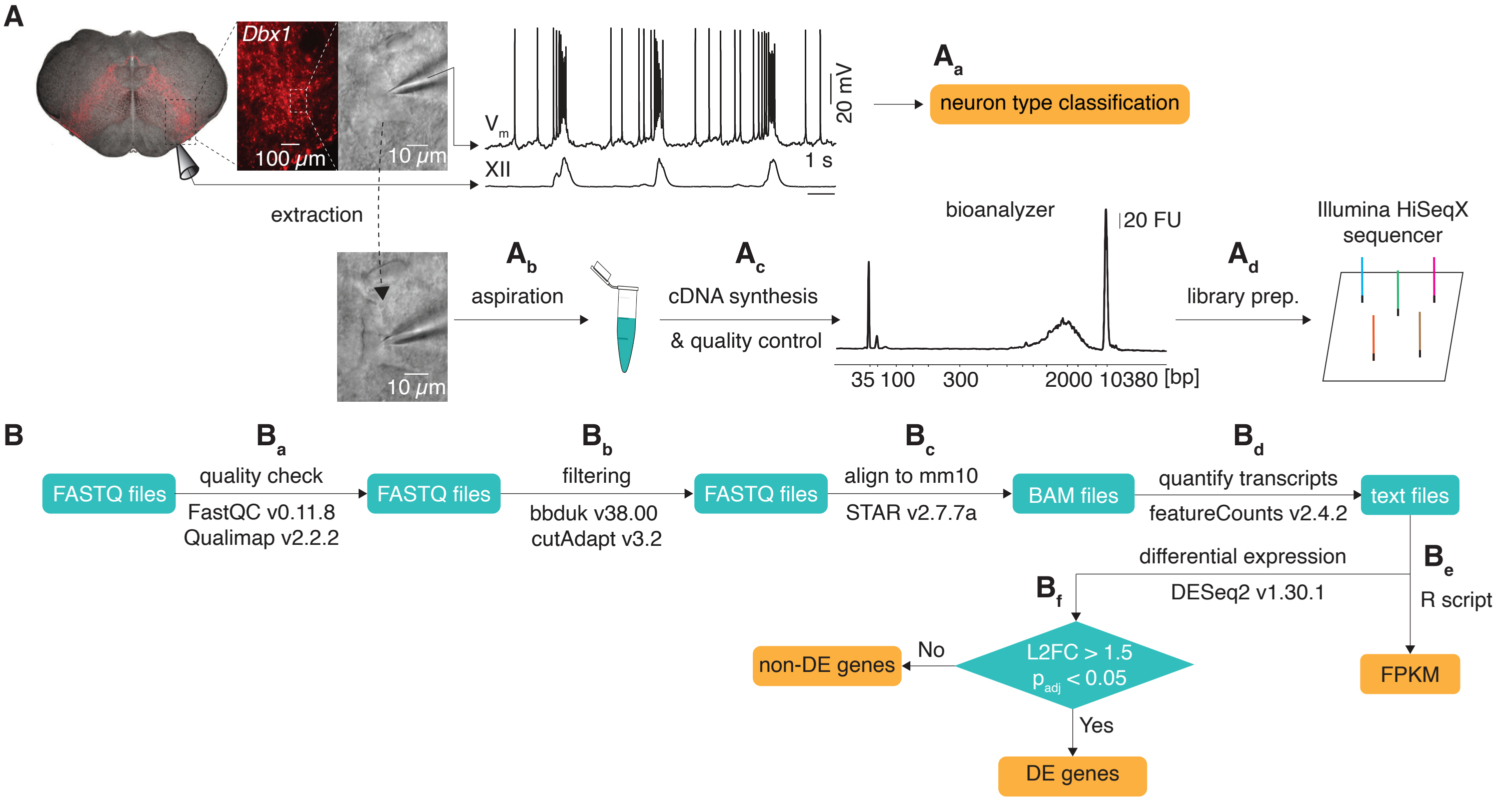


**Supplementary Figure 1.** Schematic explanation of Patch-Seq. **A,** Rhythmically active Dbx1 preBötC neurons identified by fluorescence and recorded in whole-cell conditions (Vm, top trace) with XII motor output (bottom). **$A_a$ – $A_d$** graphically represent steps in the Patch-Seq workflow as detailed in Methods. **B,** Flowchart (**$B_a$ – $B_f$**) that graphically represents the bioinformatics workflow as detailed in Methods.
