## Supplementary Fig. 2 for "Transcriptomes of Electrophysiologically Recorded Dbx1-derived Inspiratory Neurons of the preBötzinger Complex in Neonatal Mice"

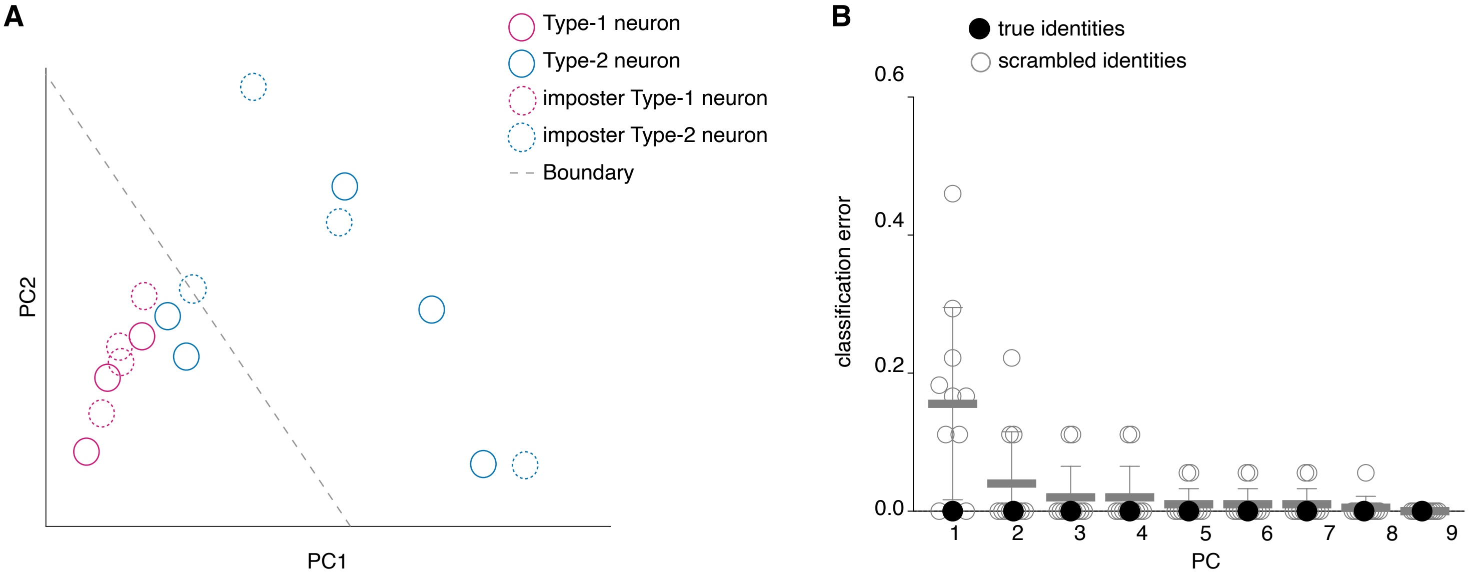


**Supplementary Figure 2.** Tests of PCA separation of Type-1 and Type-2 Dbx1 preBötC neurons. **A,** Example of PCA after scrambling the identities of 50% of the Dbx1 preBötC neurons as detailed in Methods. Neurons with intact identities are shown with solid circles. Neurons with scrambled identities are shown dotted-line circles. In both cases, magenta indicates Type-1 and blue-cyan indicates Type-2). **B,** Classification error (y-axis) for PCs 1-9 (x-axis) for the original data set of neurons whose identities have not been modified (filled circle) and for the surrogate data sets in which of 50% of the neurons are imposters with (unfilled gray circles). Relatively high classification errors only occur for groups that contain imposters, which bolsters confidence that Type-1 and Type-2 neurons are discrete classes.
