## Supplementary Fig. 3 for "Transcriptomes of Electrophysiologically Recorded Dbx1-derived Inspiratory Neurons of the preBötzinger Complex in Neonatal Mice"

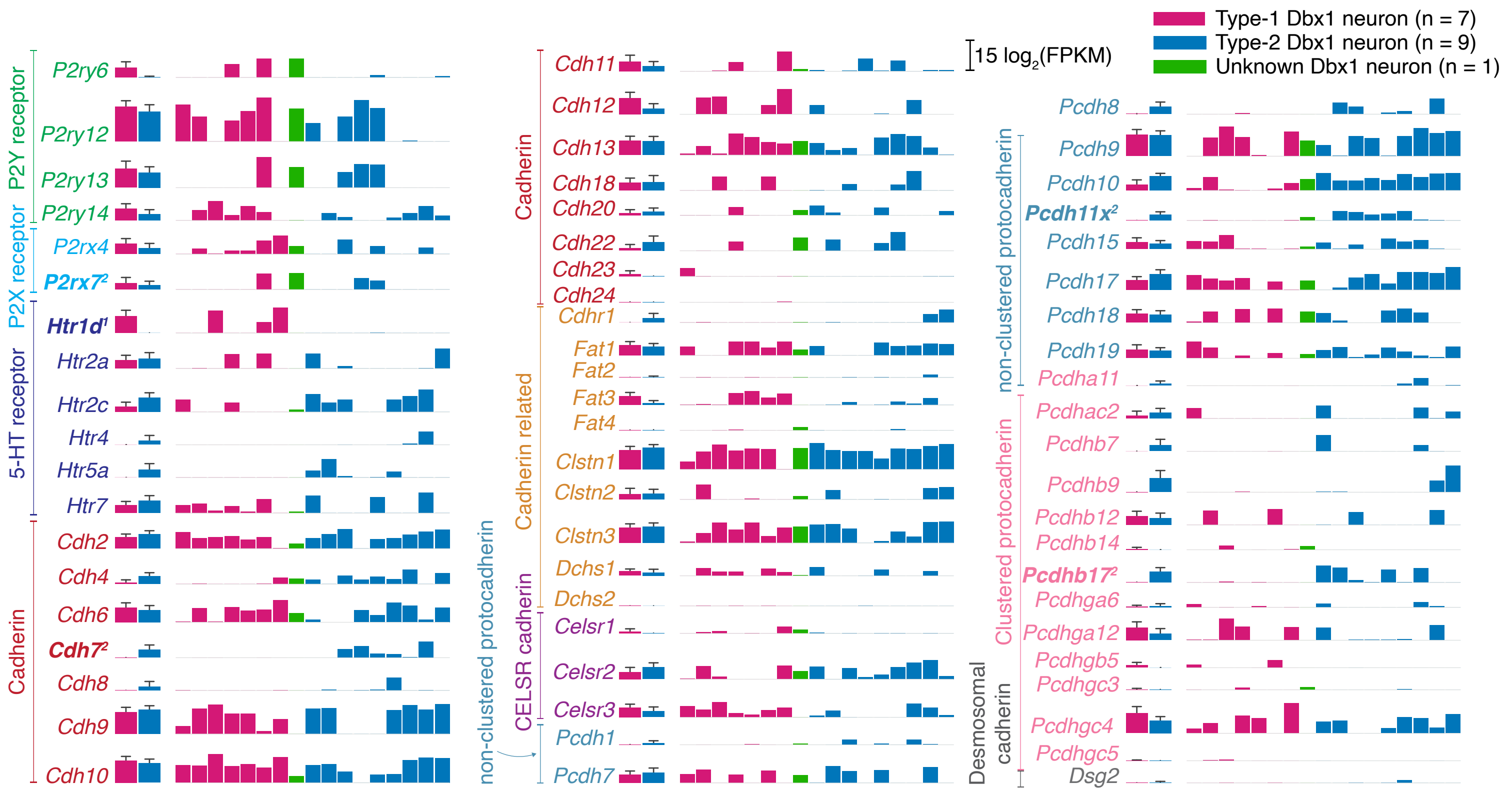


**Supplementary Figure 3.** Quantitative gene expression for purinergic receptors, serotonin receptors, and cadherins. The first two bars show group data for Type-1 (n = 7; magenta bar) and Type-2 (n = 9; blue-cyan bar). The height of the bar is $\log_2$(mean FPKM) and the error bar with horizontal cap shows $\log_2$(mean + SD). The next set of 17 bars shows $\log_2$(FPKM) values of each neuron in the following order: 7 Type-1 neurons (magenta), 1 Unknown neuron (green), and 9 Type-2 neurons (blue-cyan). Gene names are color-coded according to the subfamily to which they belong. Gene names in bold indicate DE and contain a superscript 1 if upregulated Type-1 neurons and 2 if upregulated in Type-2 neurons.
